# *Bio-Node* – Bioinformatics in the Cloud

**DOI:** 10.1101/2020.04.15.043596

**Authors:** Yannick Spreen, Maximilian Miller

## Abstract

**Motivation:** The applicability and reproducibility of bioinformatics methods and results often depend on the structure and software architecture of their development. Exponentially growing data sets require ever more optimization and performance with conventional computing capacities lacking this process. This creates a large overhead for software development in a research area which is primarily interested in solving complex biological problems rather than developing new, performant software solutions. In pure computer science, new structures in the field of web development have produced more efficient processes for container-based software solutions. The advantages of these structures have rarely been explored in a broader scientific scale. This is also the case with the trend of migrating computations from on premise resources to the cloud.

**Results:** We created *Bio-Node*, a new platform for large scale bio data analysis utilizing cloud compute resources (publicly available at https://bio-node.de). *Bio-Node* enables building complex workflows using a sophisticated web interface. We applied *Bio-Node* to implement bioinformatic workflows for rapid metagenome function annotation. We further developed “Auto-Clustering”, a workflow that automatically extracts the most suited clustering parameters for specific data types and subsequently enables to optimally segregate unknown samples of the same type. Compared to existing methods and approaches *Bio-Node* improves performance and costs of bioinformatics data analyses while providing an easier and faster development process with focus on reproducibility and reusability.

**Supplementary information** is available at https://doi.org/10.5281/zenodo.3753146

## 1 Introduction

“Free lunch is over” [1]. Moore’s law, as described by [2], is coming to an end. The exponential growth of computing performance on a single chip is dwindling. High-Performance Computing (HPC) has to scale horizontally to keep up with the explosion in data and compute requirements of new applications [1]. While computing power is no longer growing exponentially, fields like bioinformatics are generating new data at an exponential pace [3–6]. Because of this, a new challenge arises in bioinformatics computing: both data, and problem sizes grow faster than the computing power of our machines. More and more questions in bioinformatics cannot be answered by common computing approaches anymore and HPC is becoming an essential requirement. The growth in datasets gave rise to the development of multiple frameworks that enable the use of HPC resources for the computing [7–11]. However, existing solutions build a proprietary platform. Computer science research has moved on from long-winded setup processes and Docker (a containerization engine) [12] is the way forward. The many benefits of containerized deployment include reproducibility, easy setup and deployment, and a common interface for development [12]. Furthermore, the development of Kubernetes (a container orchestrator) [13] has laid the foundation for the horizontal scalability of container-deployed methods [14]. In fields like bioinformatics, computer science is a tool to solve the increasingly difficult problems of the field. Research in bioinformatics is focused mainly on the problem-solving of biological challenges rather than on the optimization of the tools at hand. This creates a discrepancy between the methods recently made available through new advances in computer science, and the approaches that are used for biological compute-heavy research. None of the common computing distribution frameworks that are used in bioinformatics were built upon existing container-based methods. [7–11] Especially the orchestration of containers in Kubernetes is not supported by any existing solution apart from *Nextflow* [10], where it is an afterthought and still an experimental feature.

Further, traditional computing methods for biological research bring tedious setup processes, which can be circumvented with the development of containers [15]. Especially since the research combines the fields of informatics and biology, many different backgrounds exist in the community and tools do not share a common approach [16]. This increases the time spent on setup and configuration for the reproducibility of results.

The benefits of deploying biological methods in containers have already been established in 2017 by *BioContainers* [17]. The growing community for this project builds a registry of thousands of containers for methods of bioinformatics. The project’s goal is to build a frame work of “*independent executable environments for bioinformatics software*” [17]. They provide a specification to standardize the submissions in the registry, but the requirements of the specification are not very strict. A format for the data inputs and outputs of a container, for example, cannot be defined [18]. This makes it hard to ensure that methods are compliant with each other when building a workflow-chain of multiple methods after another.

Even though advances are made towards container-oriented development, scaling of those methods is still left to the individual. *Galaxy*, the biggest distribution platform in bioinformatics, continues to build its proprietary ecosystem of methods and tools [9]. Building a workflow for the Galaxy project requires training to learn how to use the system since no knowledge transfer from standardized methods is possible. The *Galaxy ToolShed* [19] is a curated list of methods that are available for tasks on the platform. Custom methods cannot easily be added to a workflow because they need to be added to the curated list by the maintainers. To deploy custom methods, a whole new stack with a self-deployed webserver, database, and hosting has to be set up from scratch. A comprehensive overview of existing solutions is shown in Table 1. To overcome limitations of current solutions we developed *Bio-Node*, a universal biological computing workflow platform. *Bio-Node* aims to fulfill the needs of complex large-scale data analysis in bioinformatics while ensuring a positive user experience. To achieve this, *Bio-Node* encapsulates a complex low-level architecture with an easy to use and state of the art responsive web interface for workflow creation.

**Table 1.** Comparison of existing workflow and cloud compute solutions

|  | Slurm | Galaxy | Nextflow | CWL | Reana | Terra | BioContainers |
| --- | --- | --- | --- | --- | --- | --- | --- |
| **Pro** | Little overhead and good performance | Widely used biological analysis tool with graphical interface | Standard for workflow description and execution. | Combines Docker containers into easily reproducible and permutable workflows. | Very feature-complete system for bio data analysis with Kubernetes and CWL support. | Being built on GCP, Terra workflows can scale indefinitely by using cloud resources. | Central resource for containerized methods in biology, counting 8,000+ tools to date. |
| **Con** | Manual deployment of every job, workflows have to be done manually | Proprietary platform with barely supported self-deployment.<br>Developing custom methods is not a priority. | Textual framework without graphical components, Kubernetes support is inactive. | The standard also contains non-containerized methods and has to be implemented to build a workflow platform | No graphical interface is provided, and interactions are prone to connection issues, i.e. no chunked upload is possible. | A centralized server orchestrates all Terra interactions, making GCP the only choice for resources and self-deployment not yet supported. | No platform to run these methods is provided, workflows have to be managed by the user. |

## 2 Methods

We defined the requirements of *Bio-Node* to be a combination of Ease of Use, Reproducibility, Scalability and Data Management.

### 2.1 Ease of Use

Bioinformaticians come from various backgrounds, not necessarily involving advanced computer science and the accompanying customs. For this reason, *Bio-Node* was developed to be easy to use with as little programming and technical knowledge as possible. A priority was, to make the running of existing workflows with both established datasets, and new, user-defined data a simple task. The focus was set on self-developed methods and easily extensible repertoire of pipeline steps, making it easy to add custom steps to any type of workflow.

### 2.2 Reproducibility

Results of workflows that ran in the past should be easily and reliably reproduced by other anyone. Docker images are hence a fundamental part of Bio-Node since they integrate all code and dependencies into a permanent container. This also adds the benefit of running many methods that have been packaged into images already.

### 2.3 Scalability

*Bio-Node* should run on as many machines as are available for a given installation. Hundreds of cores should be supported without reaching any limit of the system. Since Docker is part of the architecture for previous reasons, Kubernetes is an easy choice to enable this requirement. With Kubernetes gaining traction in the Open-Source community, it’s actively developed and improved upon in recent years. This gives Bio-Node a good outlook for future support.

#### Data Management

Data sets in fields like bioinformatics are growing at an exponential rate. Every system for serious data analysis in computational biology needs to offer solid data management tools. Uploads and processing of hundreds of gigabytes of data have to be supported and manageable.

### 2.4 High-Level Architecture

*Bio-Node* was developed with the aim to overcome limitations of currently existing solutions for workflow and cloud computing. An overview of the high-level architecture is shown in Figure 1. Kubernetes serves as the main component to run and schedule jobs submitted to the queue. It offers high flexibility for hardware requirements of different tasks, rescheduling jobs once the required resources become available [20]. *Bio-Node* serves as an orchestrator of a Kubernetes cluster, without specific requirements for the cluster’s nature. The pod running the *Bio-Node* server requires management permissions for the cluster to be able to create and modify jobs and pods, and to cleanup after finished execution. The user interfaces with the platform in two main ways (Figure 1). The web interface connecting to the *Bio-Node* server provides the main instrument for data management and job control. Secondly, the Docker images, that all workflows are composed of, are uploaded to a container registry. No restrictions apply for the choice of a registry, apart from *Bio-Node* having read access to the specified images.

**Figure 1.**
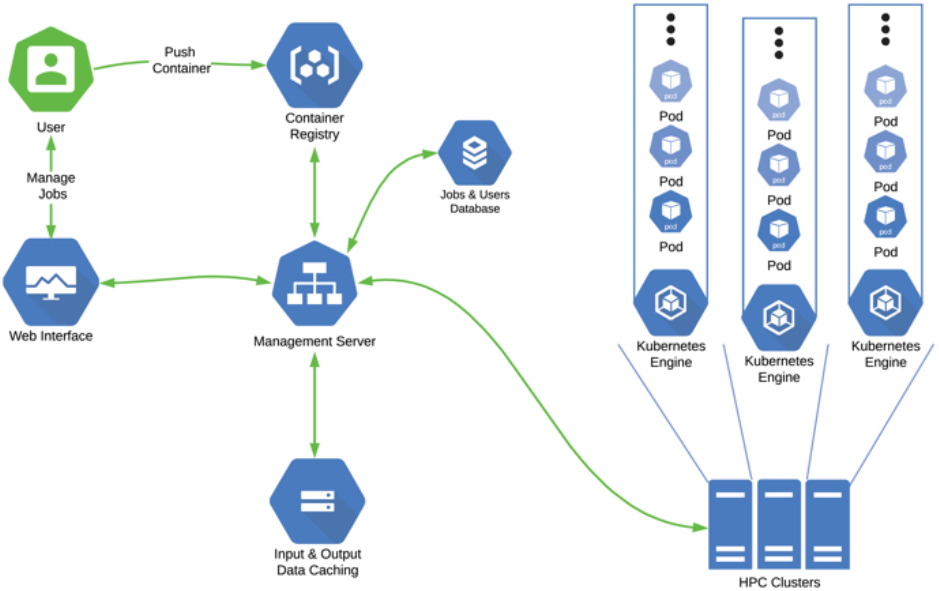
Bio-Node’s High-Level Architecture

### 2.5 Storage

One of the main requirements for a computational biology platform is a robust storage system. New data sets are released constantly and every year they grow, most of them exponentially. To work with these amounts of data, it has to be stored reliably, fast preforming and accessible. Uploads have to be especially robust so that drops in network connection do not render an entire day of uploading unusable. Regular data mounts in Kubernetes cannot be mounted with read/write permissions in more than one Docker container at a time. Concurrency and performance shortcomings require additional access management which is not internally supported by Kubernetes’ filesystem controllers. An architecture choice thus needs to be made, to address the data management needs for every Kubernetes project at hand.

For *Bio-Node*, every single step of a computing workflow is executed in its individual container. A pod, Kubernetes’ layer of abstraction for single independent Docker containers, controls its data mounts independently of the rest of the cluster. With a maximum of 100 pods per node and 10 nodes in an average target cluster, up to 1000 containers need to be able to read and write on the same volume at the same time. Among the existing storage solutions considered for *Bio-*Node (Table 2) we chose Ceph [21] for a variety of reasons. First, the Ceph distributed filesystem is built for HPC systems and supported on Kubernetes through the Rook storage orchestrator [22]. Rook allows for highly customizable deployment of storage clusters. A great range of capacities and storage performance tiers is supported. The system scales with any number of server nodes to support a high amount of parallel running mounts. All of these storage parameters are configurable for individual *Bio-Node* deployment. For our own evaluation, we chose three SSD disks with 1 TB of capacity each. At around 875 MB/s10 nominative this storage cluster is about 9 times faster than Cloud Filestore at about half the price per TB of storage. For a single job submission, we achieved 834.4 MiB/s of average throughput on four disks.

**Table 2.**
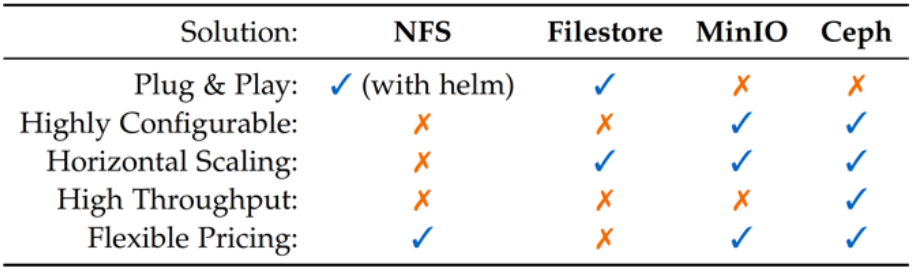
Comparison of Considered Storage Solutions

### 2.6 Auto-Clustering

We applied *Bio-Node* to implement *Auto-Clustering*, a workflow that automatically extracts the most suited clustering parameters for specific data types and subsequently enables to optimally segregate unknown samples of the same type. Clustering is an unsupervised Machine Learning (ML) technique to group data points (i.e. samples) with known and/or unknown properties into a number of close neighborhoods. If some of the samples have known labels, this method can be used for the classification of new data, where the labels are unknown. Especially in bioinformatical data analysis, many tasks [23, 24, 25, 26, 27] rely on such clustering algorithms [28, 29, 30, 31, 32]. In the first step of the actual clustering, the process is completely unsupervised, no labels are used in the analysis. One cluster is predicted for every data point, while the number of clusters in total is one of the parameters for the algorithm6. More extensive clustering frameworks, like NBClust for the R language, will optimize for the number of clusters with some automated metric. Optimizing measures like the density inside a cluster or the distance between clusters. There are many of such scoring indices that work in a completely unsupervised manner and do not require any true labels. As such, they can be seen as another parameter for the clustering algorithm. Every combination of the parameters

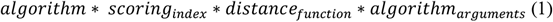

behaves very differently and performs well on different data sets. All of these options can then also be combined with different approaches at preprocessing, various dimensionality reduction methods may be used. To cluster a new data set optimally becomes a challenge. Especially if the nature of the data is not similar to preexisting sets that have been clustered in the past, many different options have to be explored and lots of manual optimization is required. We evaluated *Auto-Clustering* on metagenome functional profiles generated be mi-faser [33] for datasets made available by the CAMDA MetaSub challenges [34].

To evaluate clustering performance, we used the Adjusted Mutual Information (AMI) score [35] between the samples’ true origins and their predicted clusters is a method to assess the performance of a clustering. For two clusterings U and V, “[t]he MI measures the information that U and V share: it tells us how much knowing one of these clusterings reduces our uncertainty about the other” [35]. The order of the cluster labels is not relevant.

Here we train *Auto-Clustering* on a dataset of known samples of the 2019’s CAMDA MetaSUB challenge. We then cluster a set of unknown samples of the 2018’s CAMDA MetaSUB challenge using the set of optimal clustering parameters determined in the training step. Figure 2 shows an overview of the entire *Auto-Clustering* workflow.

**Figure 2.**
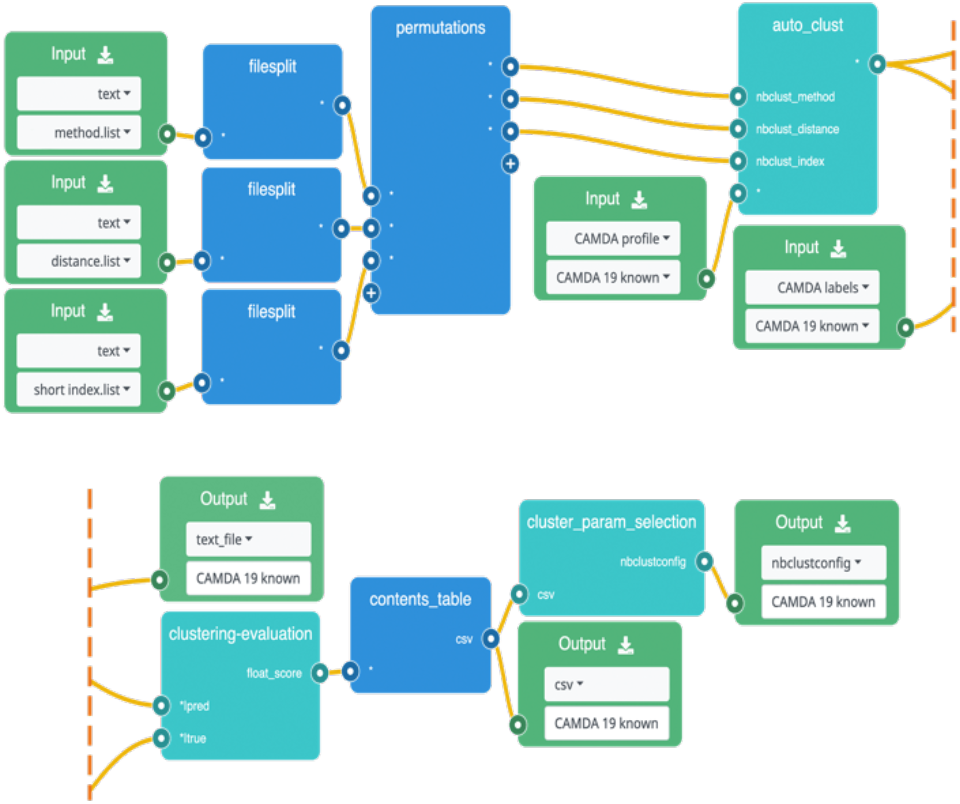
Auto-Clustering workflow

### 2.7 Performance and Cost analyses

We evaluated *Bio-Node* regarding performance and computing cost against traditional on premise HPC approaches. In detail, we compared a set of locally run compute nodes against the same configuration of nodes in the Google Cloud (18 nodes with 144 cores). We ran 100 mi-faser jobs with input datasets (metagenomic reads) ranging from 1 to 4 GB, where each job required 2 CPUs and 3 GB of RAM.

## 3 Results

### 3.1 Availability

*Bio-Node* is published on GitHub (https://github.com/bromberglab) and can be installed on any Google Cloud instance automatically using the supplied setup (see online documentation). We also maintain a publicly available *Bio-Node* instance at https://bio-node.de.

### 3.2 Performance and Cost on *Bio-Node*

We compared the compute load of three different configurations for the *Bio-Node* cluster. The system was configured to use four, eight and twelve compute nodes respectively. Every node uses eight CPU cores, resulting in a total of 32, 64 and 144 cores for each of the runs. One key architecture decision for workflows and their dependencies in *Bio-Node* is the choice to run every job for one step of the pipeline to completion before the next step is started. For the mi-faser workflow, this means that the quality control step needs to conclude for all 100 jobs of the workflow before the second step begins. This overhead is more pronounced in a cluster with low utilization because every difference in the compute time for individual jobs results in parts of the cluster being unutilized and it grows with the number of nodes. If one job takes 10% longer than all others, this job will block the entire cluster for 10% of the processing time. If only one node exists, there is no loss in compute time. With 18 nodes, however, 17 nodes wait for the last job. With longer workflow steps, more parallel jobs and higher cluster usage in general, this effect subsides. We observed a clear downward trend of total workload runtime. This downward trend is more pronounced than the upward trend in resources caused by the architectural overhead. This shows that scaling of compute resources for such analysis tasks can greatly lower the compute time and thus the waiting for analysis results. At the same time, total resource usage stays relatively constant. The resulting costs for each analysis confirm this trend. The total pricing for one workflow is nearly constant. Between $17 and $19 in GCP’s billing data, the relative difference is only 11%. At the same time, scaling up to more nodes results in a 319% speed improvement.

### 3.2 Performance and Cost on to traditional HPC

While the total runtime on *Bio-Node* for our 100 mi-faser jobs was 02:28:59, the same computations took 01:57:14 runtime on our on-premise nodes running a Slurm scheduler. This amounts to a 21.3% difference in favor of Slurm. Note that the workflow runtime was measured between the submission time of the jobs and the completion of the last job. The difference between runtime on *Bio-Node* and Slurm highlights that our architecture adds no excessive overhead to job execution times. Even though the raw runtime is about 20% longer than execution on Slurm, we established that this difference can easily be balanced by increasing compute resources and scaling up machines. However, there is a clear cost advantages which outperforms this small overhead manifold.

To analyze potential cost savings by using *Bio-Node* compared to the costs of running on-premise HPC infrastructure, we first calculate the actual runtime costs of *Bio-Node*. The 18 Node deployment we were using here, creates hourly runtime costs of USD 7.6096, resulting in USD 0.0528 per hour per core. To estimate runtime costs on Slurm we estimated over all jobs run in the last 2.5 years on our on-premise nodes. This resulted in a average load of 150,418.39 hours per core. Multiplying the load with *Bio-Node*’s runtime costs results in a total expense of USD 7,949. In comparison, the infrastructure of our on-premise nodes in a shared Slurm based HPC cluster cost USD 165,618 in total (for a limited runtime of 4 years). Being installed in May of 2017 and extended in October of 2019, the cluster has been running for 40% of its lifetime at the end of the analyzed time frame. The effective cost for this time can thus be estimated at USD 66,652. The potential cost savings thus come up to USD 58,703, or 88.1%. Bio-Node’s costs are 11.9% of those for the on-premise cluster (Figure3). We thus estimation a potential cost-saving of >80% in favor of the new cloud-based *Bio-Node* framework.

Slurm offers the baseline for scheduling overhead with very close to bare-metal performance in comparison to other systems [36]. The overhead in *Bio-Node* comprises the following steps before job execution:

1. Workflow translation to single jobs.
2. Job cross-dependency analysis and resolving.
3. Translation from jobs to single parallel sub-jobs.
4. Creation of Kubernetes resources from *Bio-Node* metadata templates.
5. Kubernetes’ handling of cluster scheduling and container creation.
6. Input processing inside the container before the tool is executed.

When a job’s execution is completed, the following steps are run afterward:

1. Event handling via Kubernetes’ web-based API in Python.
2. Output processing on *Bio-Node*’s storage volumes.
3. Dependency event handling for subsequent jobs.
4. Cleanup of temporary resources for both jobs and workflows.

Two other main factors for architectural overhead in this specific scenario are the scaling time for the cluster and the dependency semaphore for *Bio-Node* workflows. In the current version, each node in the cluster is created sequentially with around one minute of startup time per machine. A cluster with 18 nodes thus takes around 18 minutes to fully reach capacity. The second difference between the Slurm-based pipeline and running on *Bio-Node* is the synchronization of every task’s jobs. One step of the pipeline has to succeed for every input job before all outputs are handed over to the next step in the workflow. The evaluation of this architecture against raw job execution time shows, that the chosen architecture is viable. Our focus on usability does not forfeit performance and throughput significantly. It’s important to note, that the performance overhead only increases the number of core cycles required per job and not the total runtime of a workflow. The runtime is influenced by the number of nodes per cluster, impacting the compute time for a task by many magnitudes. The costs of a workflow remain constant with varying cluster sizes. We can conclude that the performance of *Bio-Node* can easily surpass all on-premise HPC clusters through GCP’s virtually unlimited scalability options. Through our cost analysis, we also show that 20% of raw compute overhead is dwarfed by the >80% cost savings.

### 3.2 Auto-Clustering performance

We explored many techniques to find an optimal clustering approach for our data at hand. Finally, a full exploration of thousands of hyperparameter combinations was the only attempt that yielded an optimal result. This exploration is a computationally intensive task and not feasible to be run on a user-machine only. Here, the use of large compute clusters saves days and weeks of runtime per data set.

Auto-Clustering in the Bio-Node framework allows us to incrementally cluster new data points through our previously learned method. Re-learning of these optimal parameters is also enabled for instances in which the dataset changes significantly. After building Auto-Clustering and using it to train on CAMDA 2019 known data samples, we evaluated our parameters with both 2018’s and 2020’s samples. We achieved a good transfer of performance for the CAMDA 2018 set. Figure 4 shows the binned results of all 1,620 parameter combinations. Clustering performances are binned with boundaries to both sides, the value named “≥ x” represents the number of clusters in between x ≤ AMI < x + 0.05. Above the threshold of the previously achieved score, only one clustering remains: the combination of the *ward.D* algorithm [37], binary distance and the *sdbw* index function [38]. This combination achieves a clustering with an AMI score of 0.585 for our data set, outperforming all other previously regarded clustering methods by a margin of 9.3% compared to our results, and a margin of 13.8% compared to our challenge submission from 2019. This result is unexpected, as it could not have been achieved without running every single permutation of the three input parameters. This combination is the only option out of over a thousand variants that outperforms our previous attempts. By automating the execution of all the jobs in parallel, we were able to improve the performance of the clustering for this data set. For 2020 however, the structure of the data was too different to achieve good results for our pre-selected configuration. Moreover, in our experience, the data was hardly cluster-able with any method. All other tested algorithms and parameters achieved AMI scores below 0.30. With the best-performing method for those samples taken into account, our pretrained parameters still performed comparatively. Since the samples are of such a different nature, however, retraining is advisable in a situation like this. For the specifics of this data set from 2020 in particular, an extension of the workflow might be advisable for the filtering of badly performing cities and analysis of the pitfalls at hand. Through the modularity of Bio-Node’s architecture and all workflows with in, these changes can be added as atomic containers very quickly. Research tasks, like the combination and trying of different tools and approaches, are accelerated through this containerized environment.

**Figure 3.**
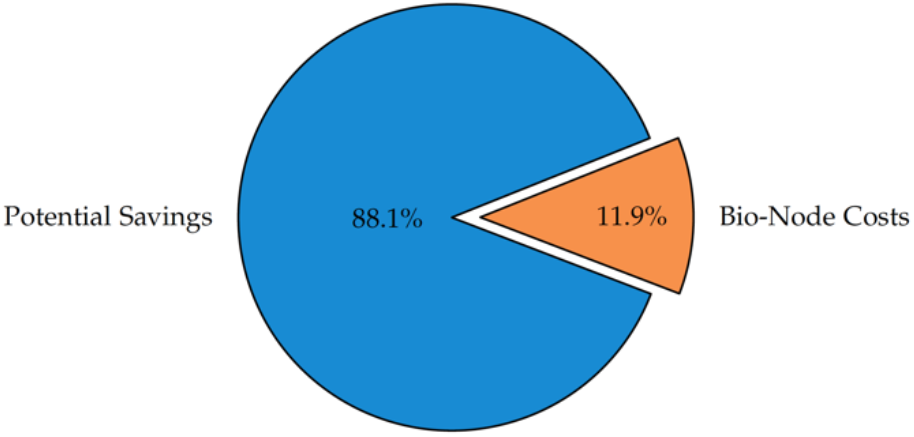
*Bio-Node* savings (USD 58,703) vs SLURM (USD 66,652)

**Figure 4.**
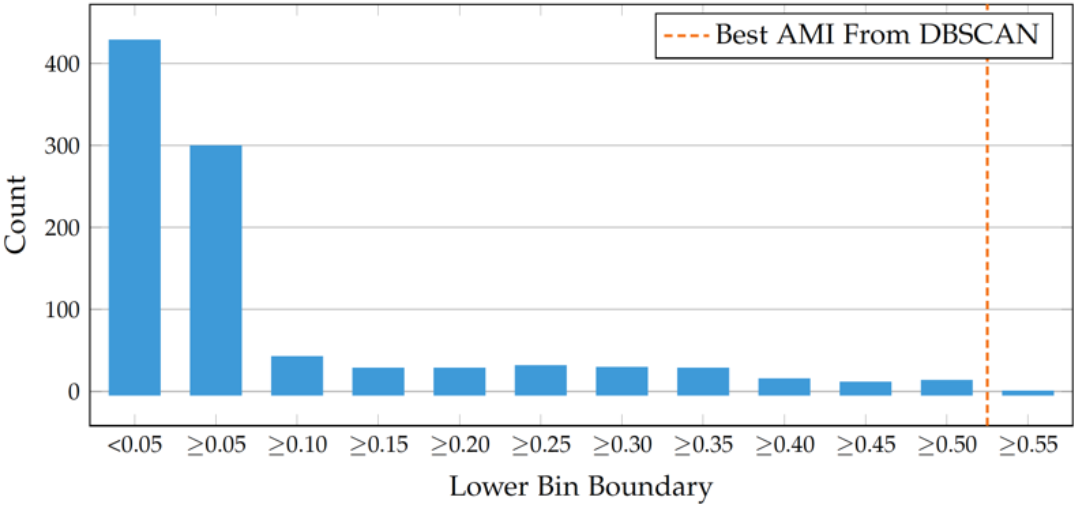
CAMDA Results with NbClust: counting the amount of parameter combinations per bin shows that only very few configurations perform well. Exactly one choice of parameters achieves a result that surpasses the best previously seen score of 0.535

As a result, we now know a parameter combination of NbClust that outperforms OPTICS and DBSCAN for the particular data set (CAMDA 2019, known) of mi-faser functional profiles for city subway metagenomes. These results were achieved without the use of any preprocessing and approaches like t-SNE and UMAP can be used to further refine them. We did outperform previous attempts with raw unprocessed data and no pre-selection of the input parameters. The alternative approaches, on the other hand, were tailored to the specifics of the given data set with weeks of optimization and research. With the Auto-Clustering results, we can now cluster future CAMDA data sets as well as unknown samples. We transferred the parameters learned from 2019’s CAMDA data to cluster the known samples from the CAMDA 2018 challenge. For this, we blinded the clustering by removing all true labels, only using them for performance evaluation. The achieved AMI score of 0.532 shows, that these data sets are similar in structure, and the difference in score is less than 10%.

## Acknowledgements

We thank the members of the Bromberg Lab (Rutgers) and Christian Dallago (TUM) for comments and valuable feedback during the development phase of *Bio-Node*.

